# Discordance between different bioinformatic methods for identifying resistance genes from short-read genomic data, with a focus on *Escherichia coli*

**DOI:** 10.1101/2021.11.03.467004

**Authors:** Timothy J Davies, Jeremy Swan, Anna E Sheppard, Hayleah Pickford, Samuel Lipworth, Manal AbuOun, Matthew Ellington, Philip W Fowler, Susan Hopkins, Katie L Hopkins, Derrick W Crook, Tim EA Peto, Muna F Anjum, A Sarah Walker, Nicole Stoesser

## Abstract

2.

Several bioinformatics genotyping algorithms are now commonly used to characterise antimicrobial resistance (AMR) gene profiles in whole genome sequencing (WGS) data, with a view to understanding AMR epidemiology and developing resistance prediction workflows using WGS in clinical settings. Accurately evaluating AMR in Enterobacterales, particularly *Escherichia coli*, is of major importance, because this is a common pathogen. However, robust comparisons of different genotyping approaches on relevant simulated and large real-life WGS datasets are lacking. Here, we used both simulated datasets and a large set of real *E. coli* WGS data (n=1818 isolates) to systematically investigate genotyping methods in greater detail.

Simulated constructs and real sequences were processed using four different bioinformatic programs (ABRicate, ARIBA, KmerResistance, and SRST2, run with the ResFinder database) and their outputs compared. For simulations tests where 3,079 AMR gene variants were inserted into random sequence constructs, KmerResistance was correct for 3,076 (99.9%) simulations, ABRicate for 3,054 (99.2%), ARIBA for 2,783 (90.4%) and SRST2 for 2,108 (68.5%). For simulations tests where two closely related gene variants were inserted into random sequence constructs, KmerResistance the correct alleles in 35,338/46,318 (76.3%) ABRicate identified in 11,842/46,318 (25.6%) of simulations, ARIBA in 1679/46,318 (3.6%), and SRST2 in 2000/46,318 (4.3%). In real data, across all methods, 1392/1818 (76%) isolates had discrepant allele calls for at least one gene.

Our evaluations revealed poor performance in scenarios that would be expected to be challenging (e.g. identification of AMR genes at <10x coverage, discriminating between closely related AMR gene sequences), but also identified systematic sequence classification (i.e. naming) errors even in straightforward circumstances, which contributed to 1081/3092 (35%) errors in our most simple simulations and at least 2530/4321 (59%) discrepancies in real data. Further, many of the remaining discrepancies were likely “artefactual” with reporting cut-off differences accounting for at least 1430/4321 (33%) discrepants. Comparing outputs generated by running multiple algorithms on the same dataset can help identify and resolve these artefacts, but ideally new and more robust genotyping algorithms are needed.

**Impact statement:** Whole-genome sequencing is widely used for studying the epidemiology of antimicrobial resistance (AMR) genes in bacteria; however, there is some concern that outputs are highly dependent on the bioinformatics methods used. This work evaluates these concerns in detail by comparing four different, commonly used AMR gene typing methods using large simulated and real datasets. The results highlight performance issues for most methods in at least one of several simulated and real-life scenarios. However most discrepancies between methods were due to differential labelling of the same sequences related to the assumptions made regarding the underlying structure of the reference resistance gene database used (i.e. that resistance genes can be easily classified in well-defined groups). This study represents a major advance in quantifying and evaluating the nature of discrepancies between outputs of different AMR typing algorithms, with relevance for historic and future work using these algorithms. Some of the discrepancies can be resolved by choosing methods with fewer assumptions about the reference AMR gene database and manually resolving outputs generated using multiple programs. However, ideally new and better methods are needed.

## 4. Introduction

Whole genome sequencing (WGS) has become a major tool for characterising the epidemiology of bacterial antimicrobial resistance (AMR) genes, representing a potentially highly discriminatory, non-targeted approach with significant advantages over other more targeted molecular techniques(1). In addition, WGS-based antibiotic susceptibility prediction has been successfully implemented as part of diagnostic and treatment workflows for *Mycobacterium tuberculosis*(2). Accurate WGS-based profiling of complete AMR gene content and prediction of susceptibility phenotypes would represent an attractive option for other commonly encountered clinical bacterial pathogens, such as Enterobacterales, including *Escherichia coli*.

Several key components are required for WGS-based AMR genotyping and predictions of susceptibility phenotype, including a robust AMR gene reference catalogue linking each genetic mechanism/sequence with a given phenotype, and accurate AMR gene identification and classification algorithms. Several catalogues and bioinformatics algorithms are now available(3-9), but only limited comparative evaluation of their outputs has been undertaken. The genetic mechanisms underpinning AMR in Enterobacterales and some other bacteria (e.g. *Pseudomonas aeruginosa*) are much more complex than those in *M. tuberculosis*, and whilst some studies suggest that WGS-based genotyping holds promise for AMR gene characterisation and the prediction of antimicrobial susceptibility for several different Enterobacterales species(10-12), the limited reproducibility and reliability of such methods in a blinded, head-to-head analysis across nine bioinformatics teams has been recently highlighted(13). However, this study was small (n=10 sequencing datasets, n=7 isolates), encountered a limited set of typing discrepancies, and used highly selected samples, meaning the impact of these issues on larger, real-world datasets remains unclear.

We therefore used simulations and three large, independent and diverse *E. coli* sequencing datasets to investigate the robustness and reproducibility of four widely-used WGS-based AMR genotyping methods (ABRicate, ARIBA, KmerResistance, and SRST2) at scale, investigating any encountered discrepancies.

## 5. Methods

### AMR gene identification methods

We evaluated the impact of different bioinformatics tools using the same AMR gene catalogue, namely the ResFinder database (v.29/10/2019). At the time the study was designed (March 2018), to be included bioinformatics tools had to: (i) have publicly available code, (ii) run on local computing architecture without major modification, (iii) accept different AMR gene databases to ensure broad and long-term typing usability, and (iv) have a command line interface that could enable batch processing of large numbers of samples (**Table S1**).

We identified four publicly available bioinformatic tools that met these criteria and used distinct AMR gene identification approaches: ABRicate(14) (which searches for AMR genes in assemblies using *BLA*STn), SRST2(7) (which maps reads directly onto the formatted AMR gene database using Bowtie 2), ARIBA(6) (which combines these two approaches, first mapping reads to the AMR gene database using minimap, and then creating local assemblies of the mapped reads using Fermi-lite) and KmerResistance(8) (which analyses shared k-mers between the query sequences and reference sequences in the AMR gene database) (**Fig. S1**). To mimic broad usability, each program was run using default parameters. For ABRicate, assemblies were first produced using SPAdes(15) run with default parameters.

### Simulated data:single and multiple allele identification, and low coverage scenarios

Prior to evaluating real data, we considered the accuracy of each method in identifying known AMR gene alleles “inserted” into simulated flanking sequence constructs. For this, each AMR gene variant in the ResFinder database (n=3,079) was flanked by 1kb of random sequence (using Numpy v1.16.4(16) and combined using BioPython(17) v1.74) and reads simulated at 40x coverage using ART (details and rationale in Supplementary Methods, **Fig. 1, S2**). Other ART parameters were: error profile=“HISEQ2500”, mean DNA fragment length (standard deviation)=480bp (150bp), and read length=151bp. Each bioinformatic method was then tested to see if it could correctly identify the AMR gene variant, using default parameters. We repeated this 10 times to assess variability across repeats.

**Figure 1.**
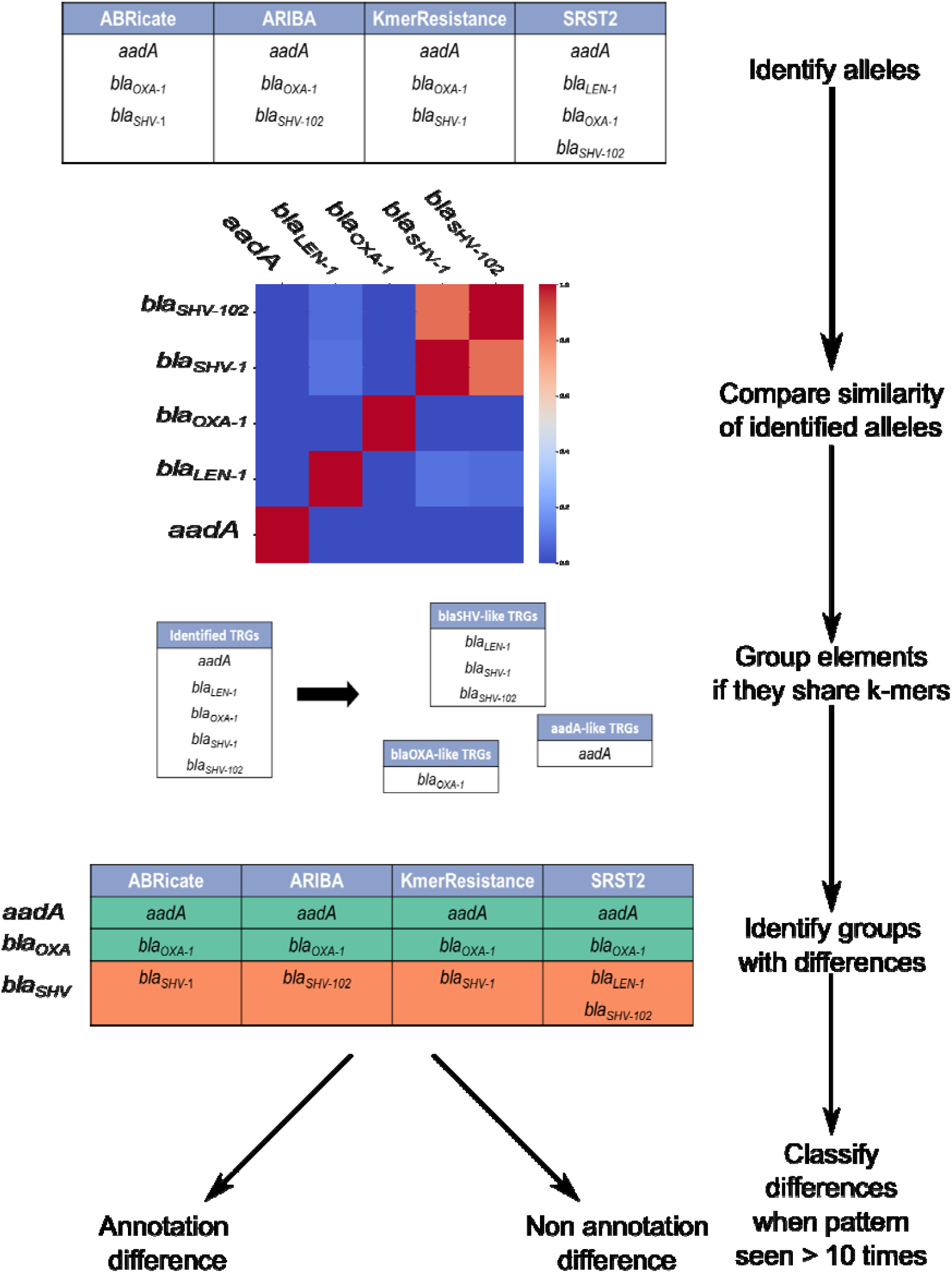
Methods to identify discrepancies.

We also considered two *a priori* scenarios that are thought to affect AMR genotyping(18), namely a *multiple allele scenario* in which multiple closely genetically related alleles (see below) of a given AMR gene were present, and a *low quality* scenario reflected by low sequencing coverage. For the *multiple allele* scenario we excluded target AMR gene variants that were incorrectly identified individually by any method (see Results), and then calculated pairwise nucleotide similarity between all remaining AMR gene variants. To do this, each remaining AMR gene variant was split into 31-mers, which were then compared with 31-mer sets from every other non-excluded AMR gene variant using pairwise Jaccard’s similarity indices. AMR gene variant pairs were defined as similar if they shared any 31-mer, resulting in a total of 46,318 possible similar AMR gene variant pairs (**Fig. S3-S5**).

For the *low coverage scenario*, reads were simulated from 176 *bla*_TEM_ gene-containing constructs at coverage depths ranging from 1x to 50x using ART (n=176*50=8,800 simulations), reflecting total *bla*_TEM_ diversity present in the ResFinder database at the time of simulation. Each construct contained a random perfect reference *bla*_TEM_ variant flanked by 1kb of random sequence on each side produced using Numpy/BioPython as above. Simulated reads were then processed by each genotyping method using default settings and the identified variants were compared with the known *bla*_TEM_ variants present in each construct. The measure of performance for this scenario was the proportion of *bla*_TEM_ variants correctly identified by each method at each coverage level.

### Real data:Isolate selection

To evaluate performance on real data, we then studied a total of 1,818 *E. coli* isolates comprising three different WGS datasets in order to reflect different strain-level and AMR gene diversity: (i) 984 sequentially collected bloodstream infection isolates at Oxford University Hospitals (OUH) NHS Foundation Trust(19) (“Oxford dataset”); (ii) 497 animal commensal *E. coli* isolates donated by the UK Animal and Plant Health Agency (APHA)(20) (“APHA dataset”), and (iii) 337 *E. coli* isolates collected by UK Health Security Agency’s (UKHSA) Antimicrobial Resistance and Healthcare Associated Infections (AMRHAI) Reference Unit, which investigates isolates enriched for rare or important resistance genotypes encountered in the UK (sequenced for this study, “UKHSA dataset”).

Isolates were re-cultured from frozen stocks stored in nutrient broth plus 10% glycerol at -80°C. DNA was extracted using the QuickGene DNA Tissue Kit S (Kurabo Industries, Japan) as per manufacturer’s instructions, with an additional mechanical lysis step (FastPrep, MP Biomedicals, USA) immediately following chemical lysis. A combination of standard Illumina and in-house protocols were used to produce multiplexed paired-end libraries, which were sequenced on an Illumina HiSeq 2500, generating 151bp paired-end reads. High quality sequences were de-novo assembled using Velvet(21) as previously described(22). *In silico* Achtman(23) multi-locus sequence types (MLST) types were defined using ARIBA(6).

While this work does not attempt to predict resistance from WGS data, each isolate had linked AST (summarized in **Table S2, Fig. S6**), which we have included as the complexity of resistance genotype identification is associated with the phenotype. Isolates had complete AST data available for: ampicillin, ceftazidime and one other 3rd generation cephalosporin (cefotaxime for the animal commensal isolates, ceftriaxone for all others), gentamicin, ciprofloxacin, and co-trimoxazole.

We compared AMR genotypes reported for each isolate by each method (**Fig. 1)**. Reported alleles were grouped into gene families using 17-mers .Discrepancies were classified according to which of the four bioinformatics methods agreed. The cause of discrepancy was investigated for all patterns occurring in at least 10 isolates. To investigate less common errors we assessed all discrepancies which affected *bla*_TEM_ alleles.

## 6. Results

### Simulated scenarios

#### Accurate identification of single AMR gene variants in simulated sequence constructs

For the 3,079 AMR gene variants in the ResFinder database, all four genotyping methods correctly identified those inserted into random sequence contexts for all repeats in 1999 (64.9%) cases. ARIBA was the only tool to intermittently correctly identify alleles across repeats (N=42/3079 (1.3%) alleles), being always correct for 2,783/3,079 (90.4%) alleles. All other tools were consistent across repeats, with KmerResistance being correct for 3,076 (99.9%) simulations, ABRicate for 3,054 (99.2%), and SRST2 for 2,108 (68.5%) (**Fig. 2**). For SRST2, most errors were due to its approach of pre-clustering reference sequences into sub-families by sequence identity prior to genotyping, thereby essentially excluding *a priori* the possibility of identifying alleles that were not selected as the representative for these sub-family clusters. This error is explained in more detail below as it also affected genotyping in real isolate sequences.

**Figure 2.**
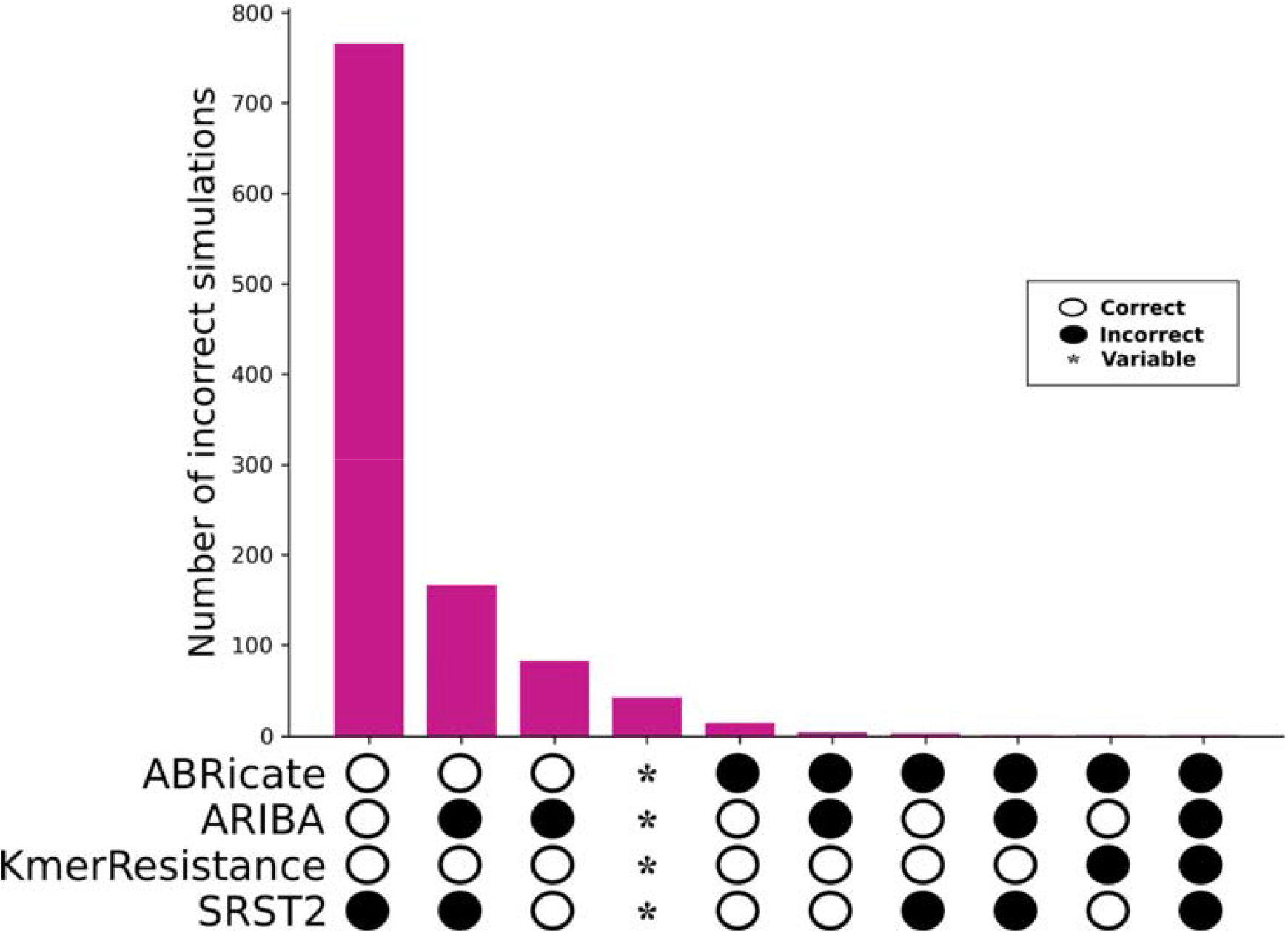
Identification of known single AMR gene variants in simulated contexts by bioinformatic method. Note only cases where one or more methods were incorrect are shown (n=1,081). * signifies where genes were variably correctly identified across 10 repeats.

### Impact of the presence of multiple closely related alleles on genotyping calls

The multiple allele simulation caused problems for all algorithms. Overall, KmerResistance was the most successful, identifying both correct genes in 35338/46318 (76%) of scenarios (**Table 1**). By contrast the next most successful was ABRicate, achieving this in 11842/46318 (26%) with the remaining tools, ARIBA and SRST2 only achieving this in 1679 (4%) and 2000 (4%) scenarios respectively. In addition, assembly-based algorithms frequently failed to completely assemble genes, with ABRicate and ARIBA reporting fragmented/incomplete alleles for 26848/58499 (46%) and 2483/18830 (13%) of the correct alleles they identified respectively. SRST2, as expected, found only a single allele in most (33319/46,318 (72%)) cases, as dictated by its clustering parameters. Unsurprisingly all four programs were most likely to make erroneous genotyping calls as the simulated pairs of alleles became more closely related (**Fig. S7a, Fig. S7b**).

**Table 1.** Performance of genotyping methods in evaluating simulated constructs with two related allelic variants. Percentage reported out of a total of 46,318 simulations performed for each method.

|  | Number of calls (%) |  |  |  |
| --- | --- | --- | --- | --- |
| Genotyping call | ABRicate | ARIBA | KmerResistance | SRST2 |
| No correct calls | 6355 (14%) | 29571 (64%) | 1795 (4%) | 10298 (22%) |
| One correct call but additional incorrect calls | 21012 (45%) | 164 (<1%) | 1028 (2%) | 184 (<1%) |
| One correct call, no incorrect calls | 415 (1%) | 14500 (31%) | 8150 (18%) | 33319 (72%) |
| Two correct calls, but additional incorrect calls | 6712 (14%) | 404 (1%) | 7 (<1%) | 517 (1%) |
| Two correct calls, no incorrect calls | 11842 (26%) | 1679 (4%) | 35338 (76%) | 2000 (4%) |

### Impact of sequencing depth on genotyping calls

KmerResistance was able to identify *bla*_TEM_ alleles at lower coverage than any of the other methods (**Fig. 3**). Above 15x depth of coverage for the gene, all methods correctly identified *bla*_TEM_ alleles in simulated constructs in > 95% of cases (**Fig. 3**). All methods were able to identify all of the *bla*_TEM_ alleles correctly at least once, but examples existed for all methods where the allele was correctly identified at low coverage, but then mis-classified at higher coverage. In general, ABRicate and SRST2, while requiring greater sequencing depth to correctly identify *bla*_TEM_ alleles initially were more accurate at higher coverage depths, making erroneous calls for only 1/176 (0.6%) and 0/176 (0%) of *bla*_TEM_ alleles at depths >20x. In contrast, for >20x coverage ARIBA and KmerResistance made erroneous allele calls for 23/176 (13%) and 6/176 (3%) *bla*_TEM_ variants respectively. Above 40x coverage ABRicate was incorrect for one (0.6%), ARIBA for four (2%), KmerResistance for one (0.6%), and SRST2 for zero (0%) simulated *bla*_TEM_ alleles.

**Figure 3.**
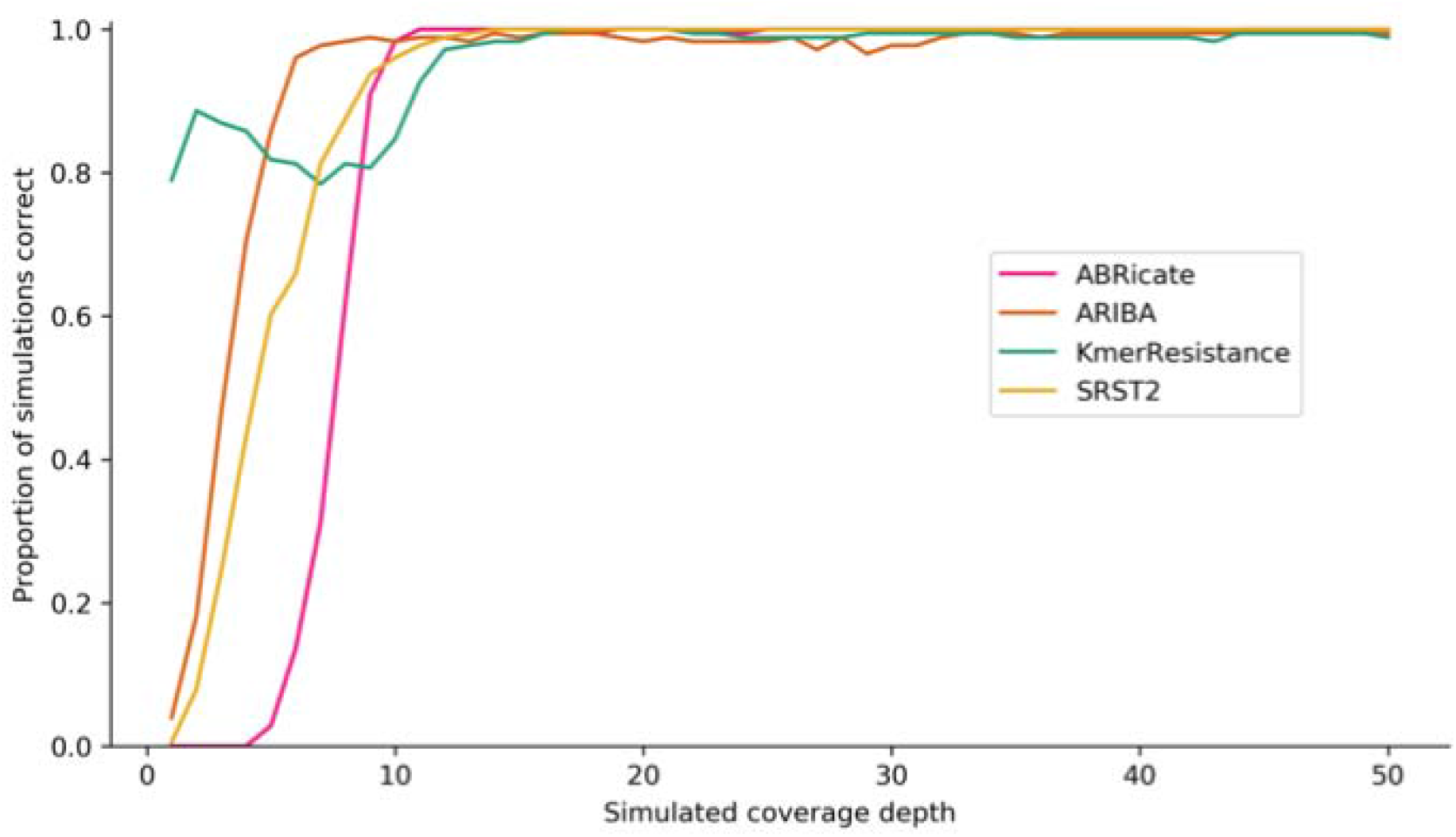
Proportion of correct genotype calls for single AMR gene variants in simulated constructs by coverage depth and bioinformatics method.

### Real data

#### E. coli isolate diversity, antimicrobial susceptibility phenotypes and antimicrobial resistance genotypes

The 1,818 isolates were diverse, representing >260 multi-locus sequence types (STs), which were differentially distributed among the datasets. For example, although ST131 was the most common (207/1818 (11%) isolates), this was largely due to the fact it was by far the most common in the UKHSA dataset (74/337 (22%) isolates). In the Oxford dataset, it was only the second most common ST (123/984 (13%) isolates) after ST73 (161/984 (16%)) isolates) and it was rare in the APHA isolates (10/497 isolates (2%)).

Correspondingly, the set also contained a broad range of resistance genes, but the exact number was dependant on the method of search. For legibility, we have included results as reported by ABRicate as this is the most conceptually simple and interrogatable approach.. The commonest AMR-associated sequence identified was *mdfA*. This is known to be universal in *E. coli*, and correspondingly was identified in all 1,818 isolates in the dataset. There were no other ubiquitous AMR genes; however, several were common across datasets, with *bla*_TEM_, *aad*A, *sul, tet*, and *dfr* genes occurring in >40% of the isolates. As expected, more UKHSA isolates contained extended-spectrum beta-lactamase (54/337 vs 94/1481) and carbapenemase (18/337 vs 1/1481) genes (p=<0.001). Aside from *bla*_TEM_, other beta-lactamases were rare among the APHA dataset. Outside of beta-lactam-associated AMR genes, the Oxford dataset had the lowest proportion of other AMR genes for all the different gene families encountered in this study.

### Genotyping discrepancies

10487 different genes (N=15588 different alleles) were identified in the 1818 isolates by the four methods. 1392/1818 (76%) isolates had discrepancies across the four bioinformatics methods for at least one gene. At the gene-level, aside from for *tet, aadA* and *cat* genes, the performance of the bioinformatic tools was similar (**Fig. 4, panel a**), with tools reporting each gene in the approximately same proportion of isolates (within +/-2%). With regards to the three outliers, ABRicate reported *tet* and *aadA* genes in 19% and 10% more isolates respectively than the other three tools, and ABRicate and KmerResistance reported *cat* genes in 5% more isolates than ARIBA and SRST2. By contrast, the alleles reported by each tool were often discrepant, with alleles of some genes (e.g. *blaSHV, blaCMY)* consistently being differentially reported (**Fig. 4, panel b**). Consequently, pairwise agreement between any two different tools was less than 59% (N=1065 isolates, **Fig. 4, panel c**). While unsupported genotype reports (i.e. where the output of one tool was not supported by any other) were common for all tools (**Fig. 5**), KmerResistance reported fewer unsupported genotypes than the other three tools (p<0.001).

**Figure 4.**
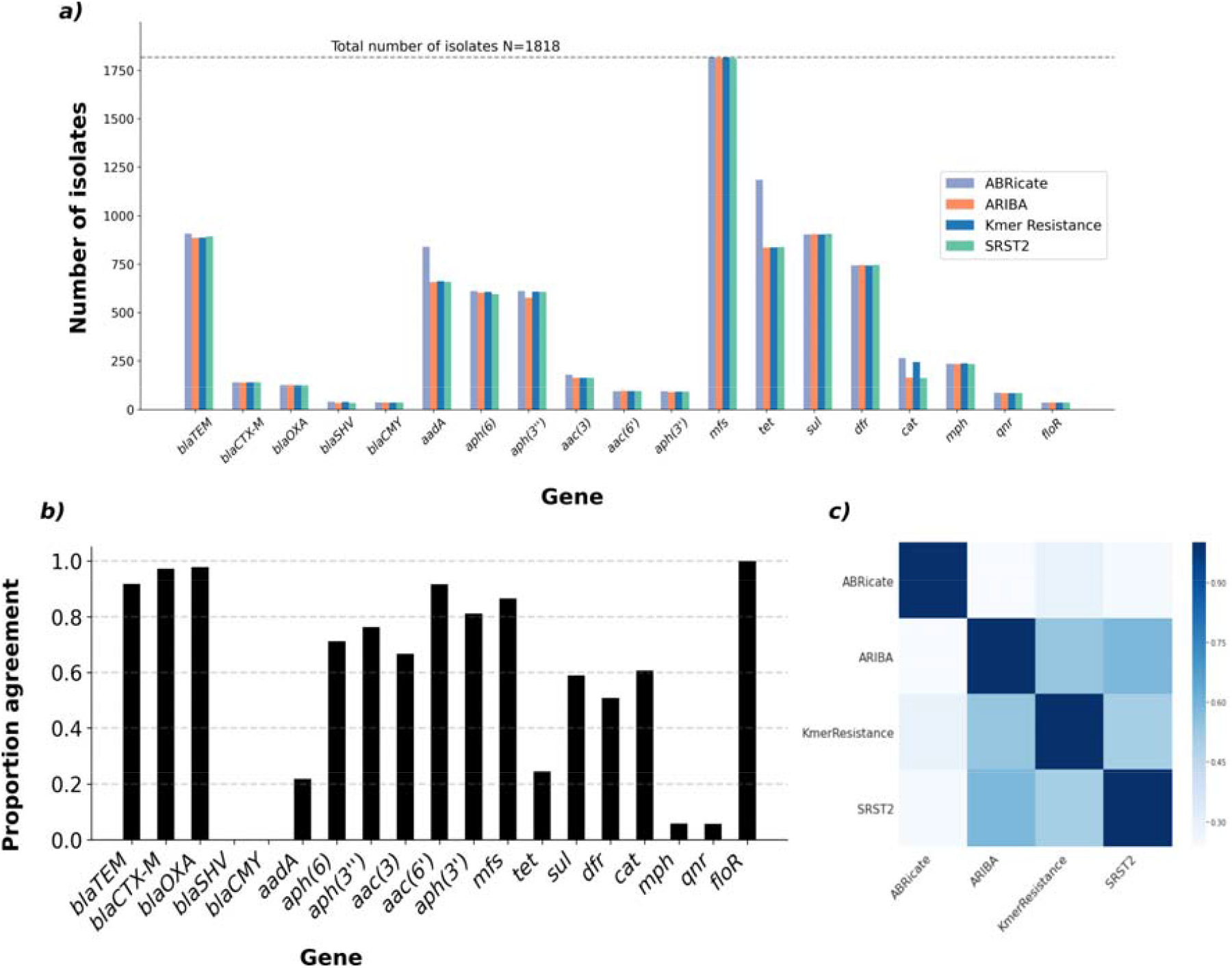
Gene identification concordance vs allele identification concordance. Note panel a) the number isolates containing at least allele of the named genes (x-axis) stratified by method. Panel b) what proportion of all times a given gene was identified did the four methods agree, e.g. for every time *bla*_SHV_ it was discordantly reported by the four bioinformatic methods. Panel c) pairwise agreement between the different methods across all isolates.

**Figure 5.**
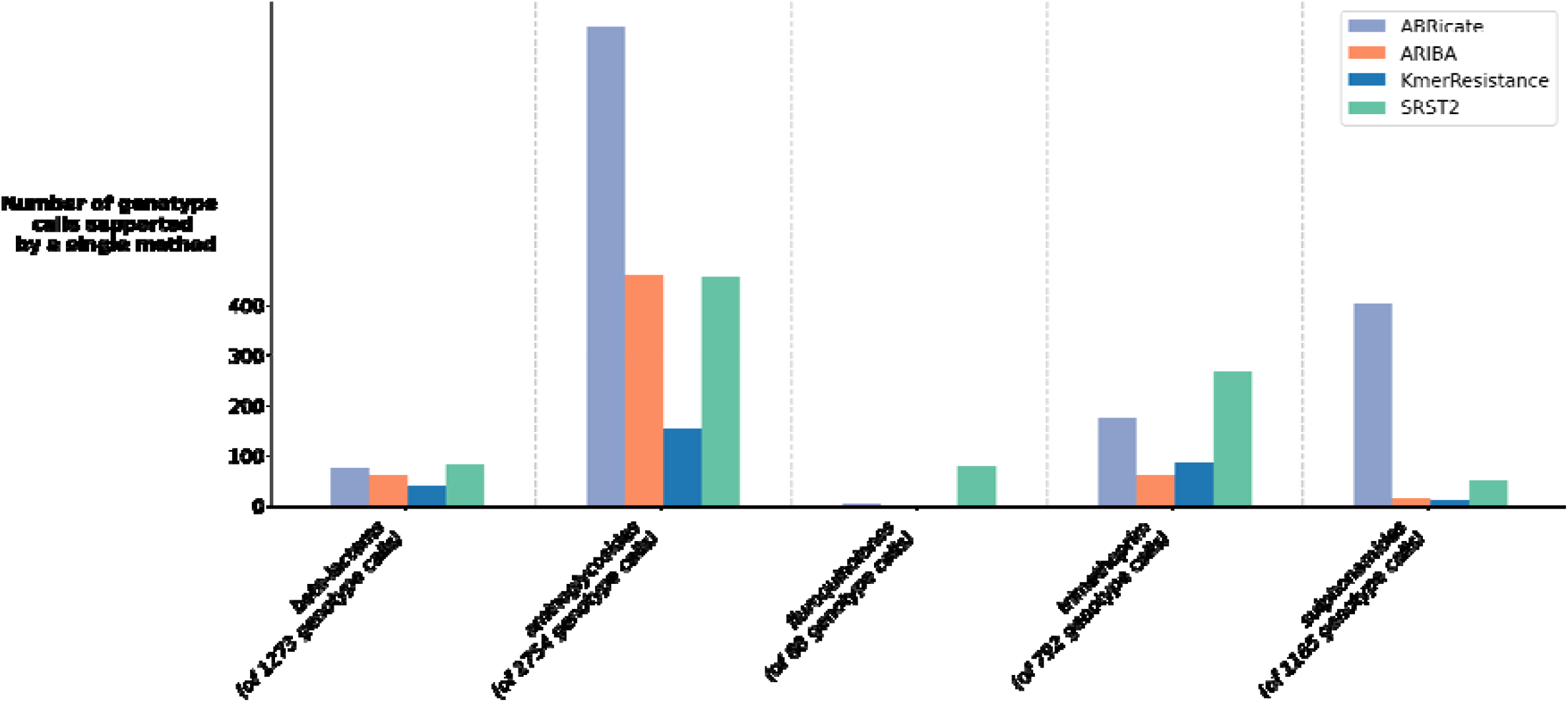
Genotype calls produced by a single method only, stratified by antibiotic class.

### Causes of genotyping discrepancy

At least 2530/4321 (59%) of allele-level discrepancies were due to programs naming the same underlying sequence differently (annotation differences). We identified three different causes of differences through investigation of discrepantly reported genes: differences due to difficulty distinguishing between optimal matches among alleles with nested sequences (N=1737 genes), spurious identification of additional alleles due to reads being multiply mapped to distant variants of the same allelic family (N=547 genes), and finally tools choosing different optimal matches based on optimal DNA sequence alignment when the database only contains one sequence per protein (N=197) (**Fig. 6**). These issues occurred alone in 1944/2530 (77%) discrepantly reported genes, and correspondingly 586/2530 (23%) had multiple issues occurring in combination. In isolation these errors typically caused only a single method to be discordant, but when combined resulted in more complex patterns of discrepancy and could make all four methods disagree with one another. In addition to annotation, ABRicate’s lesser requirement for complete gene coverage (which aims to mitigate assembly errors) caused at least 1430/4321 (33%) allele-level discrepancies. Discrepancies less easily classified as (but still possibly related to) annotation/cut-offs did occur, but thankfully only affected 381/10487 (4%) of reported genotypes.

**Figure 6.**
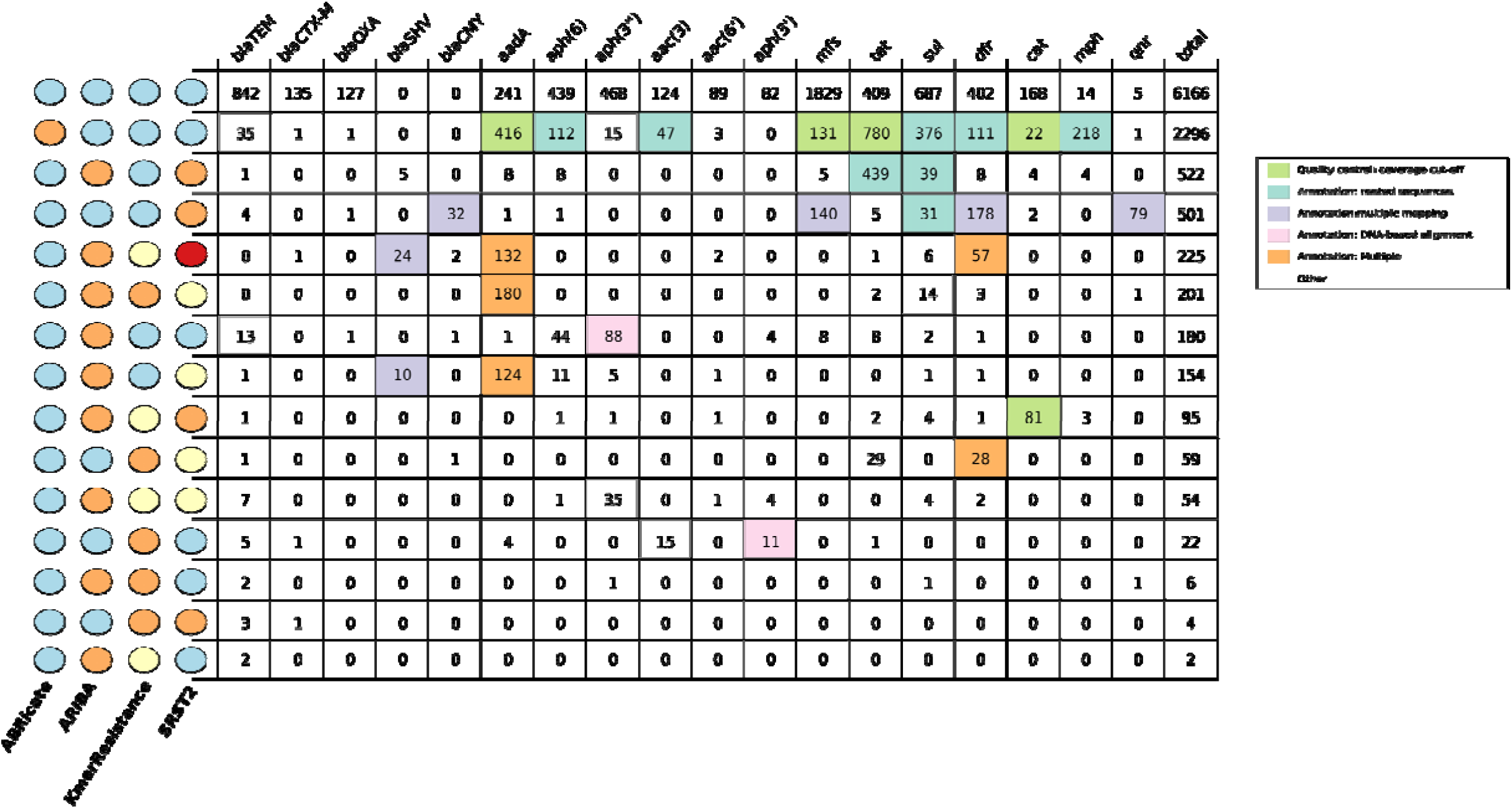
Genotyping agreement across all four bioinformatics algorithms evaluated, stratified by gene. Note colours on the left indicate which methods eed with one another, with circles the same colour indicating agreement. Colours in the main panel of the figure were used to identify the cause of the crepancy, as denoted in the figure key.. Cells (in the figure) were coloured if > 90% of isolates were caused by a given discrepancy. Cells with less 10 lates were not investigated.

### Annotation related discrepancies

The most common type of annotation error (N=1737 genes) was the result of tools struggling to choose optimal matches when the database contains nested sequences. One such example of this seen in our data caused by the sequences for two different *dfrA7* alleles in the October 2019 Resfinder database, dfrA7_1_AB161450 and dfrA7_5_AJ419170. The shorter of the two (dfrA7_1_AB161450, 474 base pairs long) aligns almost perfectly (percentage identity = 99%, 1 single nucleotide gap) with the first 473 bases of dfrA7_5_AJ419170. ARIBA, KmerResistance and SRST2, which look for the best identity sequence matches, all report the sample contains a perfect match for dfrA7_1_AB161450. By contrast ABRicate, which uses *BLA*ST to identify optimal sequences, reports the sample contains a near perfect match to dfrA7_5_AJ419170, as with this being a longer match it is more statistically significant. This then occurs each time *a dfrA7* allele is encountered, occurring 24 times in our dataset. Similar errors occur for several other genes, including *sul, tet, aph(6)*, and *aac(3)*.

The second most common annotation discrepancy (N=547 genes) was tools reporting multiple alleles due to reads mapping to two or more distant variants of the same allelic family. An example seen in our data is both ARIBA and SRST2 reporting multiple *bla*_SHV_ alleles. In this instance, ARIBA and SRST2 identify a primary perfect allele and a second allele with a lower quality match. These multiple matches however were likely spurious, with <10 reads mapping individually to each allele, no clear heterozygosity observed in read pileups, and no fragmentation in assembly graphs. This is the result of a biproduct of how mapping methods identify optimal matches. Both ARIBA and SRST2 map reads to each sequence in the database, and then compare “closely related” sequences to decide which mapping is optimal. Defining “closely related” however is not straightforward (**Fig. S8**). Reads mapping to more than one set of “closely related” sequences can result in tools finding multiple gene variants when the isolate only had one gene original

The final common annotation discrepancy (N=197 genes) was because many tools select the allele to report on the basis of which sequence in the database had the optimal DNA alignment with the target resistance gene. Resistance gene nomenclature is generally based on proteins however, and consequently resistance databases often only catalogue one sequence per resistance protein. Variant alleles with synonymous mutations fail to perfectly match any element, and may have an alternate optimal DNA match. We saw this in our data with 9 occasions where ABRicate, KmerResistance and SRST2 identified imperfect matches to aph(3”)-Ib_2_AF024602 and ARIBA identified an imperfect match to aph(3”)-Ib_4_AF313472. However, the sequence they were matching to in the SPAdes and ARIBA assembly was a 100% identity and coverage protein match to aph(3”)-Ib_5_AF321551.

### Non-annotation related discrepancies

In addition to annotation discrepancies that were caused by bioinformatics algorithms, genotyping calls were also affected by partial/low coverage of AMR gene targets and assembly fragmentation, consistent with the results from simulations. While for some of these, such as the 1430 discrepancies more definitively described cut-off related discrepancies occurring *in tet, mfs, aadA*, and *cat* genes, each program identified the same section of sequence, making it clear that the different programs had different thresholds for reporting, other situations were less clear. To investigate this in detail, we examined beta-lactamase matches which were either partial/low coverage or occurred across fragmented assemblies.

Partial/low coverage beta-lactamase genes were discrepantly found in 39 isolates (**Fig. S9**), particularly affecting *bla*_TEM_-like gene calls (29/39 cases). KmerResistance reported the presence of a beta-lactamase gene in all 39 of these discrepant cases, with calls supported to a varying degree by the other algorithms. However, in all but four cases, KmerResistance reported that the depth of the gene was less than 5x. For the four cases where the gene was present at greater than 5x depth as called by KmerResistance, three (present at depth >100x) were omitted from ARIBA reports as ARIBA assemblies contained mis-sense mutations and the final one (present at depth 17x) failed to assemble for ABRicate.

Assembly fragmentation affected ABRicate and ARIBA beta-lactam resistance gene calls in 24 cases, with 16 of these likely to be due to the presence of multiple closely related beta-lactamase alleles thereby affecting assembly integrity. The possibility of heterozygous alleles was indicated by the ARIBA flag “variants_suggest_collapsed_repeat”, and the SRST2 “minor allele frequency value” was high (>20%). KmerResistance reported two related alleles in 12/16 cases, one with high depth, percentage identity and coverage, and one much less accurately. This likely reflects KmerResistance’s winner-takes-all strategy, where matching unique k-mers to reference alleles are counted, and the reference allele with the most matches is then also assigned all reads with non-unique kmer-matches. This then leaves only reads with unique k-mers matching any closely related secondary allele, resulting in poor depth and coverage metrics.

## 7. Discussion

We evaluated the impact of bioinformatics approaches to AMR genotyping in *E. coli* for four commonly used methods and a widely used AMR gene database (ResFinder). Using >50,000 simulations and comparing >1,800 sequences sampled across human and animal reservoirs, thereby capturing common and rare AMR genotypes, we highlight that whilst currently available widely used genotyping methods are useful, their outputs should be carefully considered in light of our findings. Commonly postulated causes of discrepancy, such as low quality sequencing data, appeared to play little role. Instead, discrepancies were primarily artefactual, occurring because of different approaches in representing the complexity of the reference AMR gene database. Inconsistent labelling of gene variants will also affect the reliability of any catalogue-based methods for phenotypic prediction from WGS-based AMR genotypes. Specifically, predicting phenotype based on the presence of specific allelic variants will be clearly problematic without a reliable method of identification.

Our work agrees with previous findings by *Doyle et al*. on a small and selected dataset(13); however, we utilised large simulated and real-life datasets to identify these significant genotyping discrepancies between methods, and also characterized the underlying reasons for these discrepancies. We found most discrepancies were largely due to notation differences, i.e. each method identified the same consensus sequence but then named them differently. Further, many of these discrepancies are caused by implicit and frequently incorrect assumptions about database structure and AMR gene diversity, including that AMR genes can be classified in well-defined families using genetic identity, that different approaches to deciding best-matching alleles are equivalent, and that isolates will usually not harbor highly genetically related variants of the same AMR gene. However, nomenclature and family structure amongst AMR genes relevant to Enterobacterales is complicated, with highly diverse genotypes (and sometimes phenotypes) being assigned similar family names (e.g. *bla*_CTX-M_, *bla*_OXA_) and single SNPs in some cases leading to different resistance phenotypes (e.g. *bla*_TEM-1_ (Genbank: **AY458016.1**) - beta-lactamase inhibitor susceptible i.e. susceptible to amoxicllin-clavulanate, *bla*_TEM-30_ (Genbank: **AJ437107.1**) - beta-lactamase inhibitor resistant i.e.g resistant to amoxicillin-clavulanate). Given this, it is not surprising that we found methods that make fewer assumptions (e.g. KmerResistance) to be more robust. Based on our findings high accuracy resistance genotyping may require the use of multiple different methods to cross-check results, and/or a clear understanding of the specific assumptions underlying the methods used, before conclusions about allele presence are drawn.

One of the key strengths of this analysis was its combined use of both simulations and real world data. By using simulations, we were able to benchmark methods against a known truth, which is impossible to do with real-world data. Previous studies using only real-world data have attempted to overcome the absence of complete knowledge of the underlying genotype by using phenotypic data as a reference standard; however genotype-phenotype correlations remain poorly defined(10, 19). By subsequently using a large sequencing dataset of isolates obtained across niches, we were then able to assess the extent of discrepancies in real-life, replicating the problems observed in simulated data.

A limitation of this work is that we chose not to evaluate the impact of database choice, and this will represent future work. Currently, as has been highlighted previously(24), there are discrepancies between the AMR databases in common use, with each having a slightly different scope and in some cases differential names for different AMR gene variants (e.g.*str*A vs *aph*(*6)-Ia or aphD*, and *str*B versus *aph(6)-Id*). Comparing databases would have therefore added significant further complexity whilst limiting the generalisability of findings. A further limitation stemming from our fixed choice of database is that we have not analysed any methods where the bioinformatic method and database are intertwined (e.g. ResFinder/PointFinder or RGI). As the interaction between tool and database was the cause of many issues, it is possible that methods that are database-specific will perform better. However, the drawbacks of these combined resources are their inflexibility, again limiting generalisability. A further limitation was that these genotyping algorithms were compared using an older version of the ResFinder database – the most up to date when this work was originally planned. Since this time, 70 sequences have been added, 2 sequences modified and 2 sequences deleted (See supplementary data). We opted not to re-perform the analysis due to its manual nature and that as most of the discrepancies relate to underlying principles behind the algorithms rather than the specific implementation. Finally, we have focused our evaluation on *E. coli*, but it is likely that these issues will also more widely affect AMR genotyping, particularly of similar species with complex genotypes.

While WGS-based approaches are attractive for both characterizing AMR gene epidemiology and representing a subsequent tool for resistance prediction, this work highlights the need for caution when interpreting resistance genotypes reported by even widely used bioinformatics methods. Before WGS-based approaches can be considered reliable for use in *E. coli* (and likely other Enterobacterales), particularly for clinical decision making or replacing phenotypic data to determine epidemiological trends, database standardisation, the development of novel genotyping approaches, and improved validation and evaluation will be required.

## 8. Author statements

### 8.1 Authors and contributors

TJD, NS, AES, ASW, DWC and TEAP conceptualised the study. TD, NS, ASW, AES and MFA decided the methodology. NS, ASW, MFA, AES, DWC and TEAP supervised the project. NS, MA, MFA, MJE, KH and SH acquired and curated the data used in this study. TJD and JSW constructed software pipelines to analyse sequencing data using each of the bioinformatic tools. TJD and ASW investigated the data. TJD performed the formal analysis. NS, AES, SL, HP, AES and TEAP assisted with interpreting the cause and impact of discrepancies. TJD and NS wrote the original draft. TJD, NS, AES, PWF, TEAP and ASW assisted with data visualisation. All authors were involved in the review and editing process.

### 8.4 Ethical approval

Not applicable.

## 8.5 Acknowledgements

We are grateful to the microbiology laboratory teams at the John Radcliffe Hospital, Oxford, the Animal and Plant Health Agency, and UK Health Security Agency.

