## Supplementary Material for "Discordance between different bioinformatic methods for identifying resistance genes from short-read genomic data, with a focus on *Escherichia coli*"

**Supplementary Methods**

*Simulated data: Individual AMR allele identification*

We arbitrarily chose to simulate data at a 40x coverage value as it was above what was required for all methods to identify alleles correctly in the “low coverage” scenario (Fig.2) and (ii) it was well within the distribution of coverage of commonly observed AMR sequences (i.e. those conferring beta-lactam resistance) seen in real data (Fig.S2).

*Simulated data: ART parameters*

For all simulations, any non-depth parameters required for running ART were estimated from metrics derived from sequencing data from 100 randomly selected samples from the 1,818 real world *E. coli* sequences evaluated as part of this study.

*Simulated data: Multiple allele identification*

The choice of 31-mers to identify “similar” AMR gene alleles for each construct was based on an evaluation of several different k-mer lengths (11, 17, 21 and 31). Jaccard’s similarity as estimated using 31-mers was highly correlated with that estimated from the other k-mer sizes tested (Fig.S3).

**Supplementary Tables and Figures**

**Table S1: Bioinformatic AMR genotyping tools considered for the performance evaluation.**

| **Name** | **Method** |   **Code Public** | **Published (date of publication)** |    **Flexible database** | **Interface** | **Usable March 2018*** | **Key default parameters*** |
| --- | --- | --- | --- | --- | --- | --- | --- |
| **ABRicate** | Assembly + BLASTn | Y   | N | Y    | Command line | Y   | Percentage identity ≥ 75%  Width of coverage ≥ 0% |
| **AMRfinder** | Assembly + BLASTp/tBLASTx + Hidden Markov Models | Y | N | N | Command line | N - program not released |  |
| **ARIBA** | Mapping + local assembly | Y   | Y (2017) | Y  | Command line | Y   | Percentage identity ≥ 90%  Width of coverage ≥ 90%^†^ |
| **ARG-ANNOT** | Assembly + Blast | N   | Y (2014) | N    | Web | N - code not publicly available/web interface only |  |
| **KmerResistance** | Kmer sharing | Y | Y (2016) | Y | Command line | Y | ID threshold ≥ 70%  A species database to estimate chromosomal coverage |
| **Mykrobe Genotype** | DeBrujin Graph | Y | Y (2015) | Y | Command line | N - could only be run for *M. tuberculosis* and *S. aureus* |  |
| **Resfinder/ Pointfinder** | Assembly + BLASTn | Y | Y (2012) | N | Web +command line | N - code not publicly available/web interface only |  |
| **RGI (CARD)** | Assembly | Y | Y (2017) | N | Web +command line | N - database not flexible |  |
| **SRST2**   | Mapping | Y | Y (2014) 135 | Y | Command line | Y | Divergence ≤ 10%  Width of coverage ≥ 90% |

*: For algorithms used in the comparison.

†: ARIBA summarize requires ≥ 90% of the gene to be assembled into a single contig.

**Table S2. Summary of antimicrobial susceptibility phenotypes across the datasets for commonly evaluated antimicrobials. “**Non-susceptible” denotes isolates in either “I” or “R” categories.

|  | **APHA** | | **PHE** | | **OXFORD** | | **OVERALL** | |
| --- | --- | --- | --- | --- | --- | --- | --- | --- |
|  | **Susceptible**  **(%)** | **Non-susceptible**  **(%)** | **Susceptible**  **(%)** | **Non-susceptible**  **(%)** | **Susceptible**  **(%)** | **Non-susceptible**  **(%)** | **Susceptible**  **(%)** | **Non-susceptible**  **(%)** |
| Ampicillin | 205 (41) | 292 (59) | 74 (22) | 263 (78) | 446 (45) | 535 (55) | 725 (40) | 1090 (60) |
| 3^rd^ generation cephalosporin | 461 (93) | 26 (5) | 268 (80) | 69 (20) | 903 (92) | 81 (8) | 1632 (90) | 186 (10) |
| Ceftazidime | 444 (89) | 53 (11) | 256 (76) | 81 (24) | 909 (92) | 75 (8) | 1609 (89) | 209 (11) |
| Carbapenem | 497 (100) | 0 (0) | 328 (97) | 9 (3) | 983 (100) | 1 (0) | 1808 (99) | 10 (1) |
| Co-trimoxazole | 347 (70) | 150 (30) | 201 (60) | 136 (40) | 818 (83) | 163 (17) | 1366 (75) | 449 (25) |
| Ciprofloxacin | 218 (44) | 273 (55) | 177 (53) | 160 (47) | 707 (72) | 273 (28) | 1102 (61) | 712 (39) |
| Gentamicin | 450 (91) | 47 (9) | 266 (79) | 71 (21) | 912 (93) | 70 (7) | 1628 (90) | 188 (10) |

**Figure S1. Schematic of four different bioinformatics algorithms evaluated**

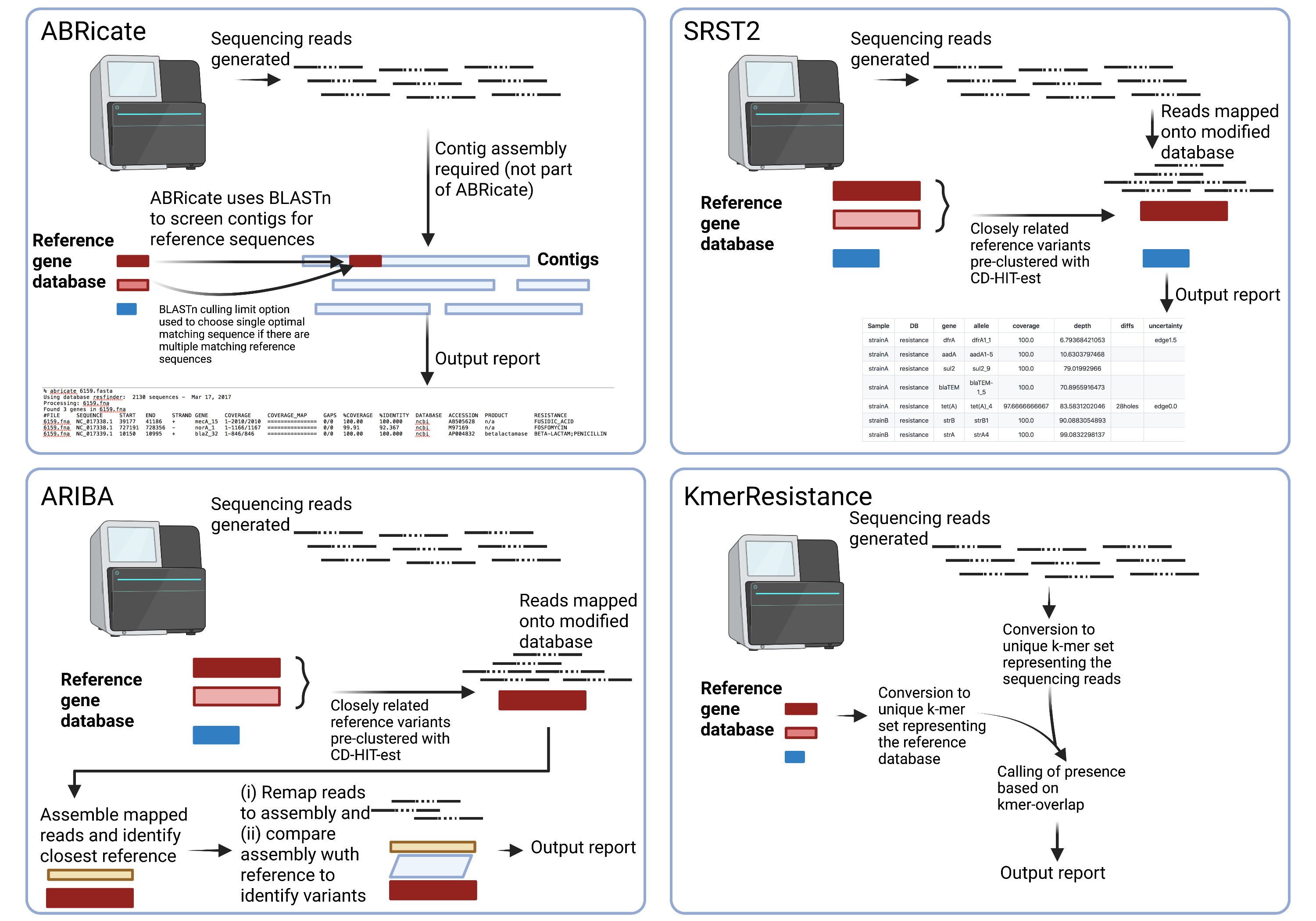

**Figure S2. Distribution of coverage depth for beta-lactam AMR genes in 1,818 real *E. coli* isolate sequences.**

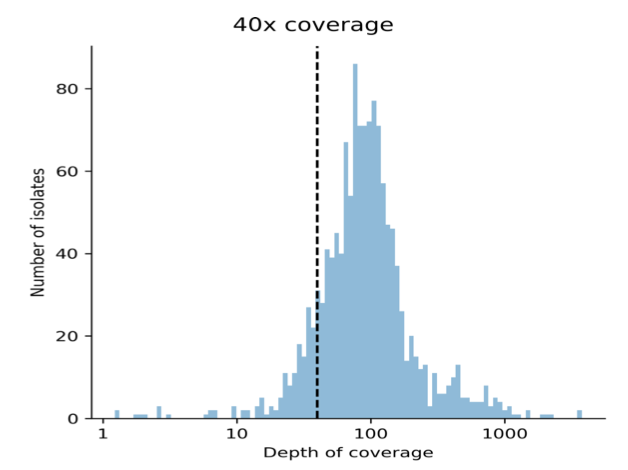

**Figure S3. Association between Jaccard index derived from 31-mers from AMR gene variants vs 11-mers (top panel), 17-mers (middle panel) and 21-mers (bottom panel).** Note that only AMR gene alleles that were correctly identified by all four bioinformatic methods were included in the multiple allele simulations (n=2,011 sequences included in the comparison).

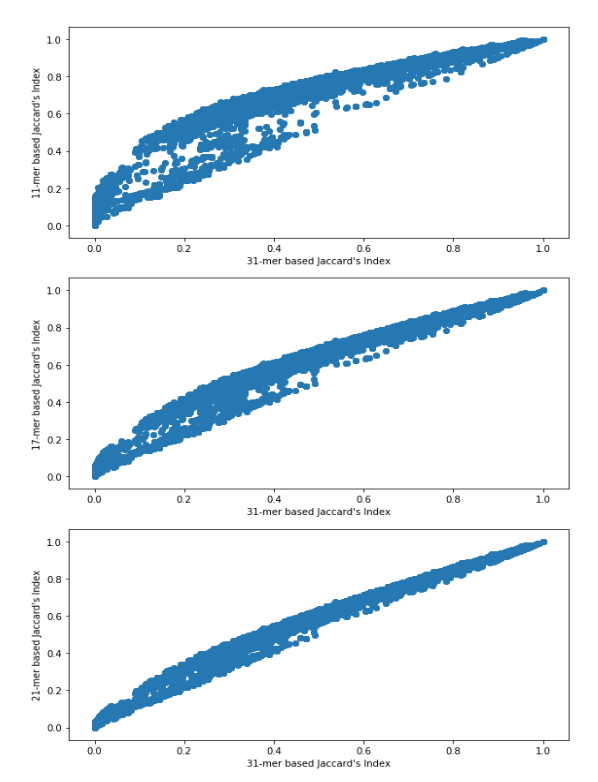

**Figure S4. Correlation of 31-mer-based Jaccard similarity index for ResFinder AMR gene allele pairs and percentage identity derived using the Needleman-Wunsch algorithm.** (n=2,011 sequences included in the comparison).

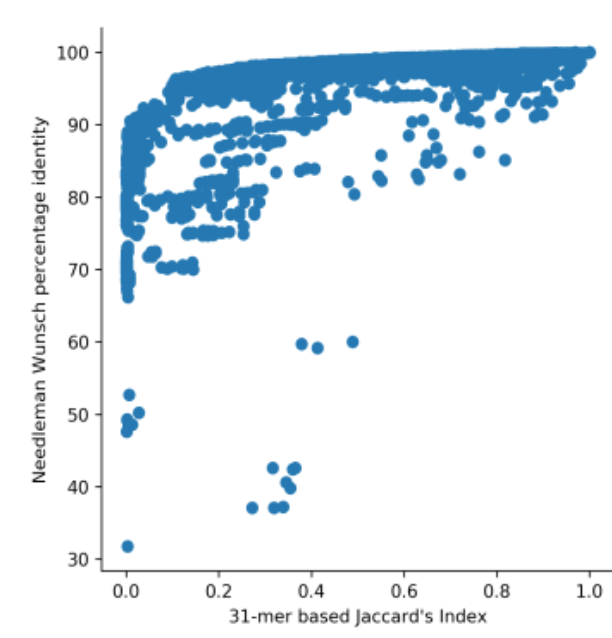

**Figure S5. Distribution of Jaccard similarity indices for ResFinder AMR gene variant sequence pairs characterized as similar based on 31-mer sharing (see Methods).**

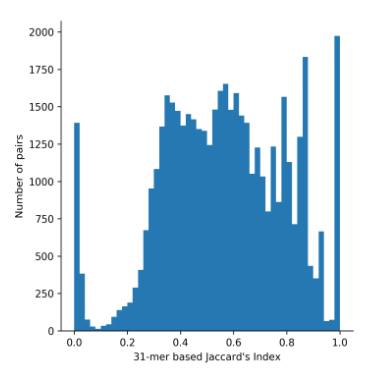

**Figure S6. MIC distributions for common antimicrobials for isolates evaluated in this study.**

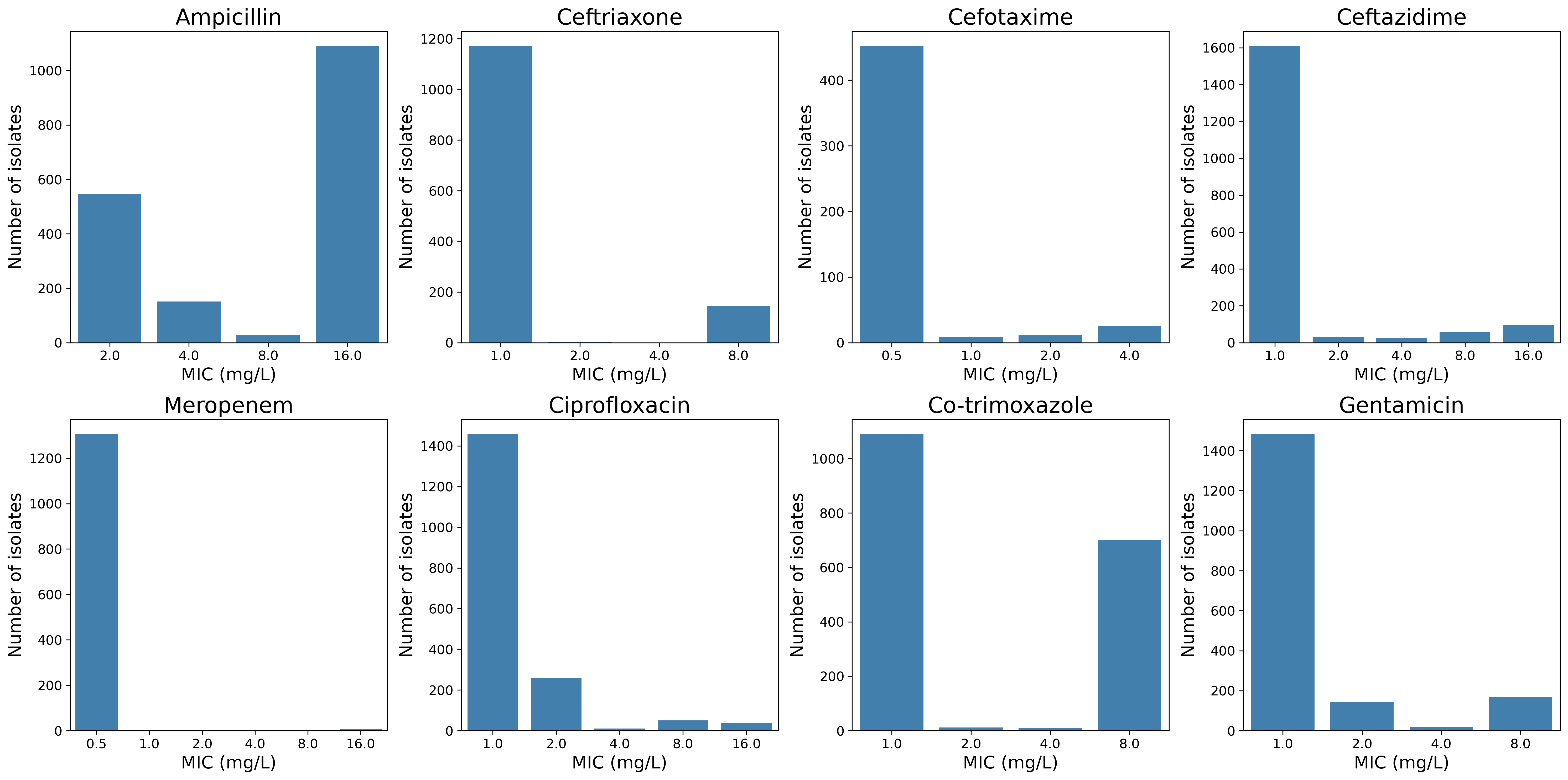

**Figure S7a. Genotyping calls for multiple allele simulations by sequence similarity of the pairs (ABRicate)**

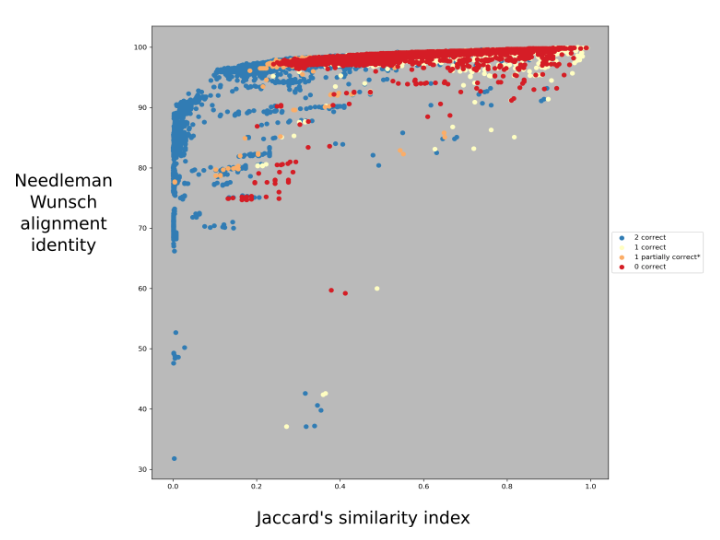

**Figure S7b. Genotyping calls for multiple allele simulations by sequence similarity of the pairs (all programs, Jaccard’s similarity)**

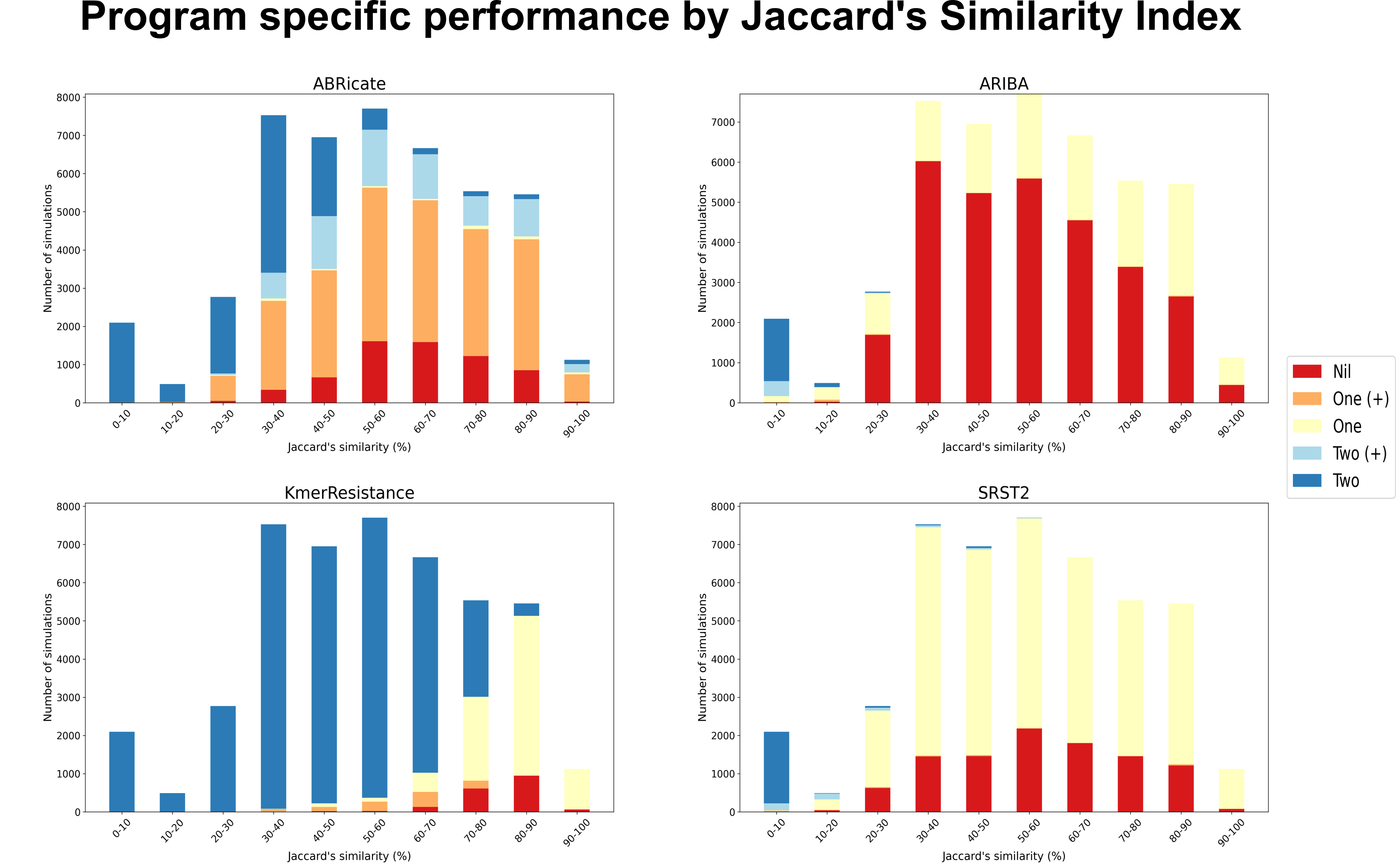

**Figure S8. SRST2 cluster size for common beta-lactamase gene families by CD-hit nucleotide percentage identity cluster threshold.** NB - x-axis is on a log scale.

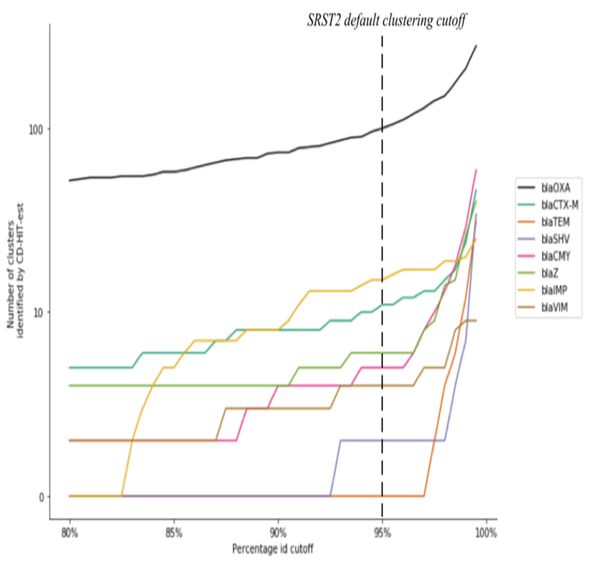

**Figure S9. Allele assignation discrepancies by method for beta-lactam resistance genes**

**Ab = ABRicate, Ar = ARIBA, Km = KmerResistance and Sr = SRST2. Discrepancy pattern indicated by left panel with white indicating gene found by this method, grey indicating not found. E.g. the first line represents 23 isolates which KmerResistance only identified as containing partial/low coverage genes.**

| Discrepancy pattern (grey indicates not found) | | | | All isolates | | Only isolates with no other beta-lactamase present | |
| --- | --- | --- | --- | --- | --- | --- | --- |
| Ab | **Ar** | **Km** | **Sr** | **Number isolates** | **Number ampicillin resistant** | **Number isolates** | **Number ampicillin resistant** |
|  |  |  |  | 23 | 6 | 17 | 3 |
|  |  |  |  | 9 | 4 | 7 | 2 |
|  |  |  |  | 3 | 1 | 2 | 0 |
|  |  |  |  | 2 | 2 | 1 | 1 |
|  |  |  |  | 2 | 0 | 2 | 0 |
| Total | | | | 39 | 13 | 29 | 6 |
